# Metagenomic analysis of marsupial gut microbiomes provides a genetic basis for the low methane economy

**DOI:** 10.1101/2024.12.04.626884

**Authors:** Kate L. Bowerman, Yang Lu, Harley McRae, James G. Volmer, Julian Zaugg, Phillip B. Pope, Philip Hugenholtz, Chris Greening, Mark Morrison, Rochelle M. Soo, Paul N. Evans

**Affiliations:** Australian Centre for Ecogenomics, School of Chemistry and Molecular Biosciences, The University of Queensland, Brisbane, Australia; Water Innovation and Smart Environment Laboratory, School of Civil and Environmental Engineering, Faculty of Engineering, Queensland University of Technology, Brisbane, Australia; Frazer Institute, Faculty of Medicine, The University of Queensland, Brisbane, Australia; Centre for Microbiome Research, School of Biomedical Sciences, Queensland University of Technology, Brisbane, Australia; Faculty of Chemistry, Biotechnology and Food Science, Norwegian University of Life Sciences, Ås, Norway; Department of Microbiology, Biomedicine Discovery Institute, Monash University, Clayton, Australia

## Abstract

The potent greenhouse gas methane is an end-product of plant biomass digestion by gut microbiota, though the amount produced and/or released varies among herbivorous animals. On a per unit of feed basis, macropodid marsupials (e.g. kangaroos) are widely thought to be low methane-emitting herbivores compared to high methane-producing ruminant livestock. How the gut microbiome contributes to the low methane status of marsupials is not well understood but of high potential value for a low methane economy. Here, we analyse the faecal metagenomes of 14 different marsupial species and 1,394 derived metagenome-assembled genomes (MAGs), focusing on the functional distinction of the bacterial and archaeal communities compared to ruminant faecal microbiomes. Though composition and function of the marsupial gut microbiome considerably varied across and within animal species, there was a clear host-associated bacterial signature for the community that differed significantly between marsupial hosts and compared to ruminants. Of particular note was a range of *Bacteroidota, Campylobacterota, Desulfobacterota*, *Pseudomonadota* and *Verrucomicrobiota* species that were enriched in marsupials and encode H_2_-uptake hydrogenases that mediate hydrogenotrophic respiration. Additionally, in support of an enrichment of electron sinks, enzymes for butyrate, propionate, and glutamate production, as well as nitrate, nitrite, and fumarate respiration were enriched in marsupials. Collectively, these data suggest that, by favoring an enrichment of alternate hydrogen sinks of bacterial origin, the low methane phenotype reported for marsupials is feasible and offers a genetic basis to pursue reductions of livestock methane emissions.

## Introduction

Atmospheric methane levels are estimated to have doubled since pre-industrial times and account for approximately 30% of global warming [1]. The largest anthropogenic source of methane is farmed ruminant livestock, such as cattle, sheep, and goats, through the action of gastrointestinal (gut) microorganisms during enteric digestion of plant biomass [2]. Gut bacteria hydrolyse and ferment carbohydrates in feed into absorbable short-chain fatty acids (e.g. acetate) and gaseous products such as carbon dioxide and hydrogen [3]. In the process of methanogenesis, methanogenic archaea oxidise hydrogen to reduce carbon compounds to methane, and in turn maintain low partial pressures of hydrogen that ensures enteric fermentation remains favourable [4]. Some of these methanogens have adapted to prevailing conditions of animal gastrointestinal tracts and are only observed in these environments based on gene surveys [5]. Different methanogen species reduce different carbon compounds: carbon dioxide for *Methanobrevibacter* species [6], organic compounds for *Methanomassilicoccales* and *Methanosphaera* species [7, 8], and a combination of the two for *Methanocorpusculum* species [9]. Much of the efforts to reduce methane emissions from ruminant livestock have largely focused on the direct inhibition of methanogens, with varying success [10–15]. One side-effect of inhibiting methanogenesis is the accumulation of hydrogen, which in turn imposes a metabolic burden on the rumen microbiota that impacts fermentation, microbial growth and generation of volatile fatty acids [16]. Thus, while partial solutions are becoming available, there remains a need for additional or alternative methods of ruminant methane reduction that draw upon a more comprehensive understanding of hydrogen flow in a broad range of animal hosts with varying digestive parameters and microbiome dynamics.

Foregut-fermenting herbivorous marsupials, such as kangaroos and wallabies (macropodids), emit lower levels of methane compared to ruminants [17–19]. This has been confirmed quantitatively with macropodids producing four- to ten-fold lower methane compared to ruminant livestock per unit of equivalent feed [20]. Methanogenic archaea are present at low abundance in the macropodid foregut [17, 19, 21] and in faecal samples from hindgut-fermenting marsupials including koalas, wombats, gliders, and possums [21, 22]. Species present include members of the genera *Methanocorpusculum*, *Methanobrevibacter* [21, 22], *Methanosphaera* [23], and *Methanomassilicoccus* [17], similar to those found in ruminants, suggesting a potential for similar methane production. However, *Methanocorpusculum* genomes show differences in alcohol dehydrogenase enzymes, both between marsupials and to ruminants, indicating alternative metabolic capabilities [22]. Limited studies also suggest differences in the wider gut microbial community and the bacterial to archaeal fraction between marsupial and ruminant species [21, 22]. Despite these recent studies, the metabolic properties of archaea in marsupials remain largely unknown and low methane emissions suggest adaptations to the marsupial gut not seen in ruminants.

Lower methane emissions by marsupials likely reflect differences in both gut anatomy and microbial activities. The shorter gastrointestinal tract and lower retention time of digesta relative to ruminants result in decreased hydrogen levels that limit methane production [3, 24]. In addition, marsupials are enriched with fermentative bacteria that produce succinate and/or propionate as an electron sink without producing hydrogen, for example with large populations of *Succinivibrionaceae* associated with reduced methane emissions in the Tammar wallaby [25]. Similar shifts towards succinate, lactate, and propionate production have been associated with low methane-emitting cattle and sheep [26–30]. Conversely, reduced methane production can also reflect the increased activity of hydrogen-consuming bacteria that reduce the electron pool available for methanogens. Indeed, recent studies have suggested ruminants harbour novel lineages of acetogenic bacteria and various hydrogenotrophic respiratory microbes that use electron acceptors such as fumarate, nitrate, and sulfate [16, 30–33]. Acetogens have also been characterised from wallaby guts, though the hydrogen sinks of marsupials have yet to be holistically examined [34, 35]. Together these results suggest a need to understand the relationship between methanogens and the broader hydrogen-cycling microbial communities of marsupials, with a systematic approach being needed to understand these differences across marsupials, not just in kangaroos and wallabies. In this study, we use genome-resolved metagenomic analyses of the faecal microbiomes from 14 diverse marsupial species to resolve the key pathways and mediators of hydrogen cycling and methane formation in these animals, and in comparison to ruminant livestock.

## Results

### Marsupial species harbour diverse and distinct microbial faecal communities

The bacterial and archaeal communities of 14 Australian marsupial species from eight marsupial families were profiled through metagenomic sequencing of 33 faecal samples (**Fig. 1, Table S1**). In addition to short-read sequencing of all samples, long-read sequencing was completed on a subset of five, selected for their identified carriage of archaeal species of interest and sequenced using adaptive sampling intended to enrich for target species. The dataset produced 1,394 metagenome-assembled genomes (MAGs) ≥50% complete with ≤5% contamination, 1,105 of these solely from short-read data and 289 from the hybrid assembly of short- and long-read data (**Table S2**). These MAGs, in addition to 1,100 public genomes estimated to represent taxa within our samples, were incorporated into a dereplicated genome database of 1,584 sequences (95% average nucleotide identity) (**Fig. S1**). Mapping of metagenomic sequence data to this database recruited an average of ∼54% of sample reads (min. 11%, max. 82%), indicating the sequencing and assembly effort captured a large proportion of the community, though some species were underrepresented (e.g. greater glider) resulting in poor discrimination between some hosts using principal component analysis (**Table S1**, **Fig. S2A-D**). Up to 78% of the unrecruited sample reads (range 17-78%, average 51%) could be aligned to contigs not included in study MAGs (**Fig. S3**). Bacterial species lacking genome representation comprise a substantial fraction of the unclassified data in some hosts, including in the greater glider, where ∼60% of unclassified reads mapped to bacterial contigs (**Fig. S3**). Non-prokaryote DNA was estimated to account for between 22% and 54% of total sample DNA based on marker gene assessment (**Table S1**), suggesting enrichment of viruses and eukaryotic microbes in some samples, however we found limited representation of this fraction within the assembled data (**Fig. S3**).

**Fig. 1.**
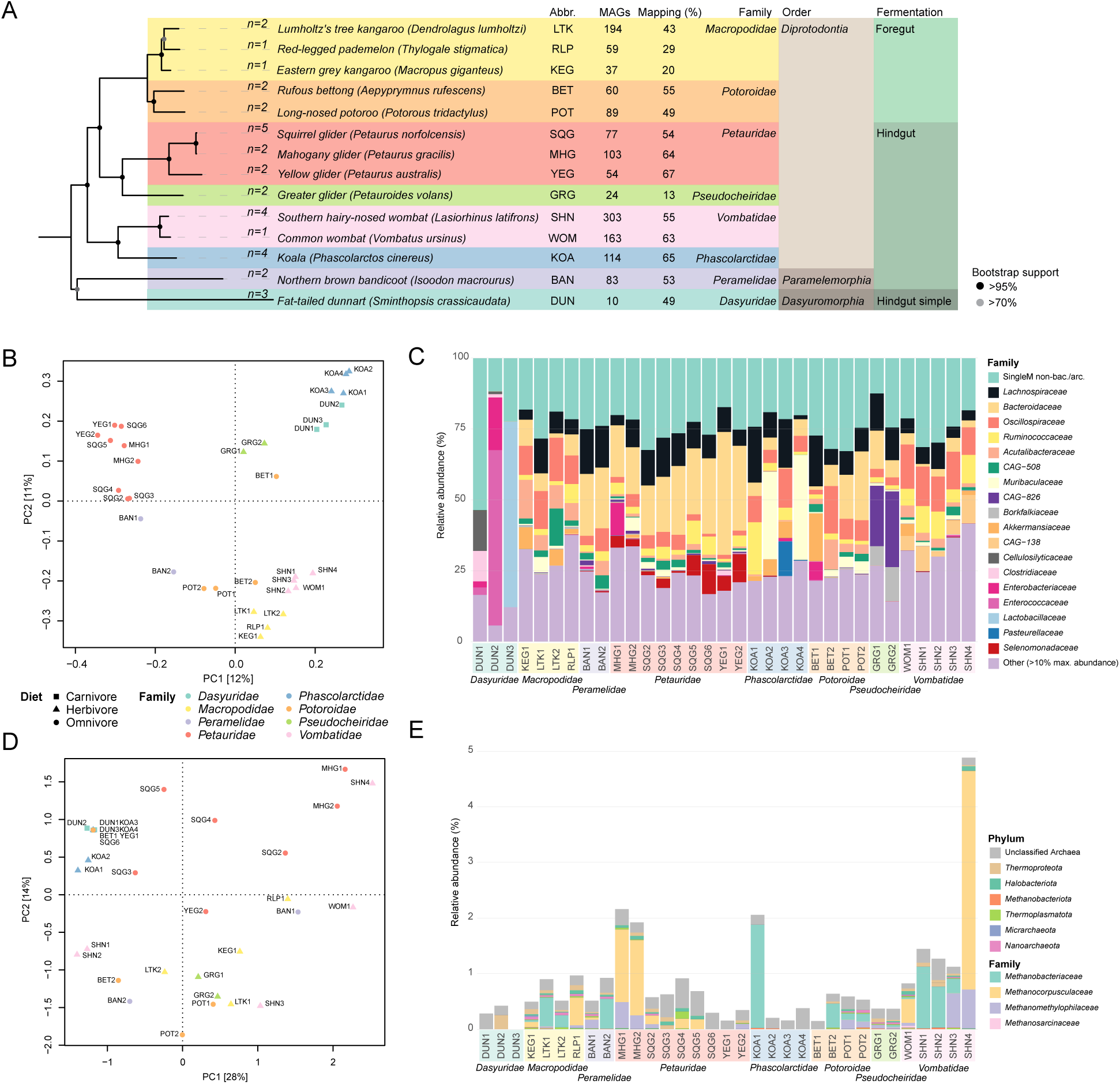
Marsupial host tree & marker gene-based community profiles. A: Maximum likelihood tree of marsupial hosts included in the dataset based on alignment of two mitochondrial (CYTB, ND2) and three nuclear (IRBP, BRCA1, vWF) genes. Bootstrap support generated from 100 replicates. Sample numbers per host indicated at tips. Abbr: abbreviated host names used throughout. MAGs: Number of metagenome-assembled genomes per host. Twenty-four genomes were recovered from faecal samples of two co-inhabiting species and are not included in host-level totals. Mapping (%): average percentage of sample reads mapping to the dereplicated genome database per host. B: PCA of faecal bacterial community. C: Bacterial and archaeal faecal community relative abundance at family level based on marker gene-based profiles. D: PCA of faecal archaeal community. E: Archaeal community family level relative abundance based on marker gene-based profiles.

Due to the incomplete representation of the faecal prokaryote community within some hosts using the genome database, the marsupial microbial communities were compared using SingleM [36]. These data support a diet-associated distinction of the bacterial community, with arboreal omnivores separated from terrestrial omnivores, herbivores and more specialised feeders (**Fig. 1B**) driven by *Prevotella*, *Candidatus* Faecousia and *Muribaculaceae CAG-873* respectively (**Fig. S4**). The only carnivorous species within the dataset, the fat-tailed dunnarts, carry distinct communities, both in comparison to other marsupials and to each other (**Fig. 1C**). Similarly, koala samples were also notably distinct from other marsupials and to each other (**Fig. 1C)**. By contrast, the archaeal community varied between individual animals and no clear separation in association with sample metadata was seen (**Fig. 1D & E**). The bacterial dominance in driving microbial variation was also evident when considering species incidence (**Fig. S5**). Redundancy analysis supported the confounded variables of host species (R^2^: 39%) and family (37%), gut anatomy (foregut/hindgut/simple hindgut) (9%), and diet (omnivore/herbivore/carnivore) (12%) as contributing significantly to bacterial community-based sample divergence when each was considered in isolation (all *p* < 0.001). No significant variables were identified for the archaeal community profiles.

We compared the SingleM-classified taxa differentially enriched in foregut and hindgut fermenting marsupials, as well as between marsupials and ruminants, in both cases excluding dunnarts. While macropods and potoroids are anatomically different to ruminants, they are foregut fermenters, making their comparison to ruminants of particular interest [17, 18]. Taxa enriched in the foregut animals are predominantly members of the class *Clostridia* including members of the orders *Christensenellales*, *Lachnospirales* and *Oscillospirales* (**Table S3, Fig. S6A**), with no taxa enriched in the more diverse hindgut fermenting marsupials. Sixty-six taxa were also differentially enriched in marsupials compared to ruminants from the family *Bovidae* (**Table S4, Fig. S6B**). Many of these taxa are known hydrogenotrophs, including fumarate and nitrate reducers (*Bilophila*, *Escherichia*) and sulfate reducers (*Bilophila*, *Desulfovibrio*), and others encode hydrogen-producing fermentative hydrogenases based on annotation of hydrogen metabolising genes within the enriched genera present in our genome database (*Phascolarctobacterium_A*, *Duodenibacillus*) (**Fig. 2**). Species of the enriched genus *Duodenibacillus* have previously been shown to be enriched in ruminants following methanogenesis inhibition [37]. Moreover, in support of our previous work, succinate-producing *Succinivibrio* and *Bacteroides* were also enriched in marsupials compared to ruminants (**Table S4**) [25]. Differential species were predominantly bacterial, with the only discriminatory archaeal species, *Enteromethanobrevibacter sp900769095* (previously *Methanobrevibacter_A sp900769095* [38]), being present in ∼15% of marsupial samples (**Table S5**). Most bacterial taxa enriched in foregut-fermenting marsupials (vs hindgut-fermenting) were identified as ruminant-associated in this analysis, supporting some overlap between these communities. Notably, the community of two of the foregut-fermenting marsupials, the red-legged pademelon and the Eastern grey kangaroo, more closely resembled the ruminant samples based on sPLS-DA (**Fig. S6B**), suggesting either taxonomic convergence or transfer of gut microorganisms between marsupials and ruminants.

**Fig. 2.**
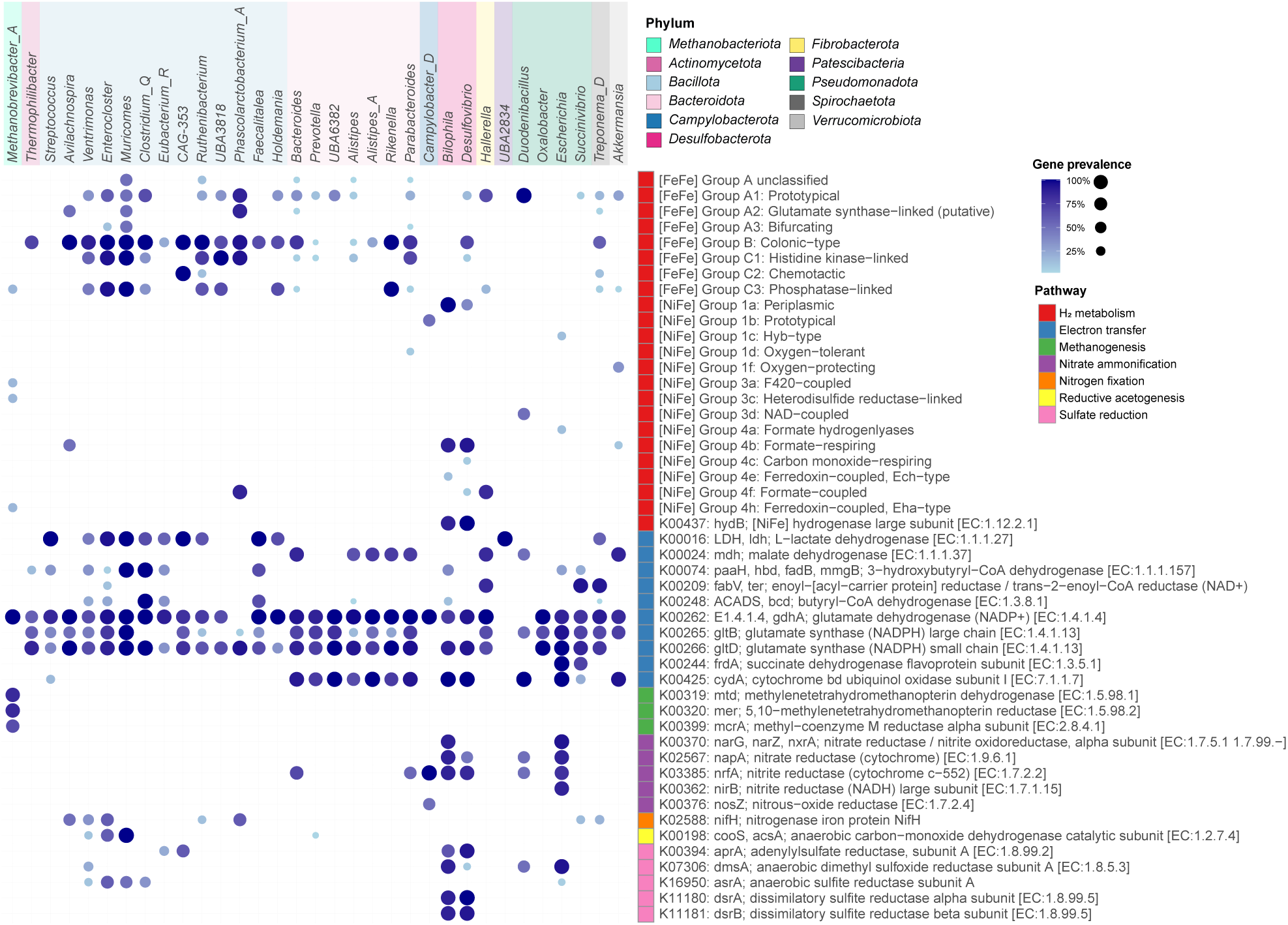
Hydrogen cycling functions across marsupial-enriched taxa. Genomes from genera within marsupi-al enriched taxa classified to genus level were functionally annotated with selected hydrogen cycling functions [32, 39]. Prevalence of each function within each genus is indicated by colour and dot size.

## Distinct carbohydrate and hydrogen-cycling pathways in marsupials

To determine whether the taxonomic variance observed between marsupials and ruminants translates to functional variance, we compiled a gene database from metagenome assemblies of all samples. Comparison of the carbohydrate-active enzyme (CAZyme) profiles between marsupials and ruminants revealed enrichment of pectin degradation capacity in marsupials, while ruminants carried broader starch-degrading capacity and an enrichment in carbohydrate-binding modules (**Fig. S7, Table S6**). Pectins are often methylated and their removal by esterases plausibly contributes to methanol availability for methylotrophic methanogens. Between marsupials, this analysis identified enrichment of host glycan-degrading enzymes in arboreal animals, driven by the *Petauridae* glider species, and increased diversity of enriched plant degradation enzymes and binding modules in the direction of predominantly terrestrial animals (**Fig. S7B, Table S7**).

To assess hydrogen flow and the potential influence of alternative hydrogen sinks on methane production in marsupials, we next used our gene database to compare the abundance of hydrogen-metabolising and reductant disposal genes between marsupials and ruminants [32, 39]. Bacterial H_2_-uptake hydrogenases involved in hydrogenotrophic respiration (group 1 [NiFe]-hydrogenases) were enriched in marsupials, while ruminants were enriched in methanogenesis-linked hydrogenases (groups 3a, 3c, and 4h [NiFe]-hydrogenases) and key methanogenesis marker genes (including *mtd*, *mer* and *mcrA*) (**Fig. 3, Table S8, Fig. S8**). Multiple electron transfer genes were enriched in marsupials, including enzymes involved in butyrate (*paaH*, *bcd*), propionate (*frdA*, *mdh*) and glutamate (*gltB*, *gltD*, *gdhA*) metabolism, supporting increased operation of these reductant disposal pathways in marsupial hosts. Marsupials were also enriched in nitrate (*narG*, *napA*) and nitrite (*nrfA*, *nirS*) reductases, supporting increased nitrate ammonification, in potential competition with methanogenesis for hydrogen.

**Fig. 3.**
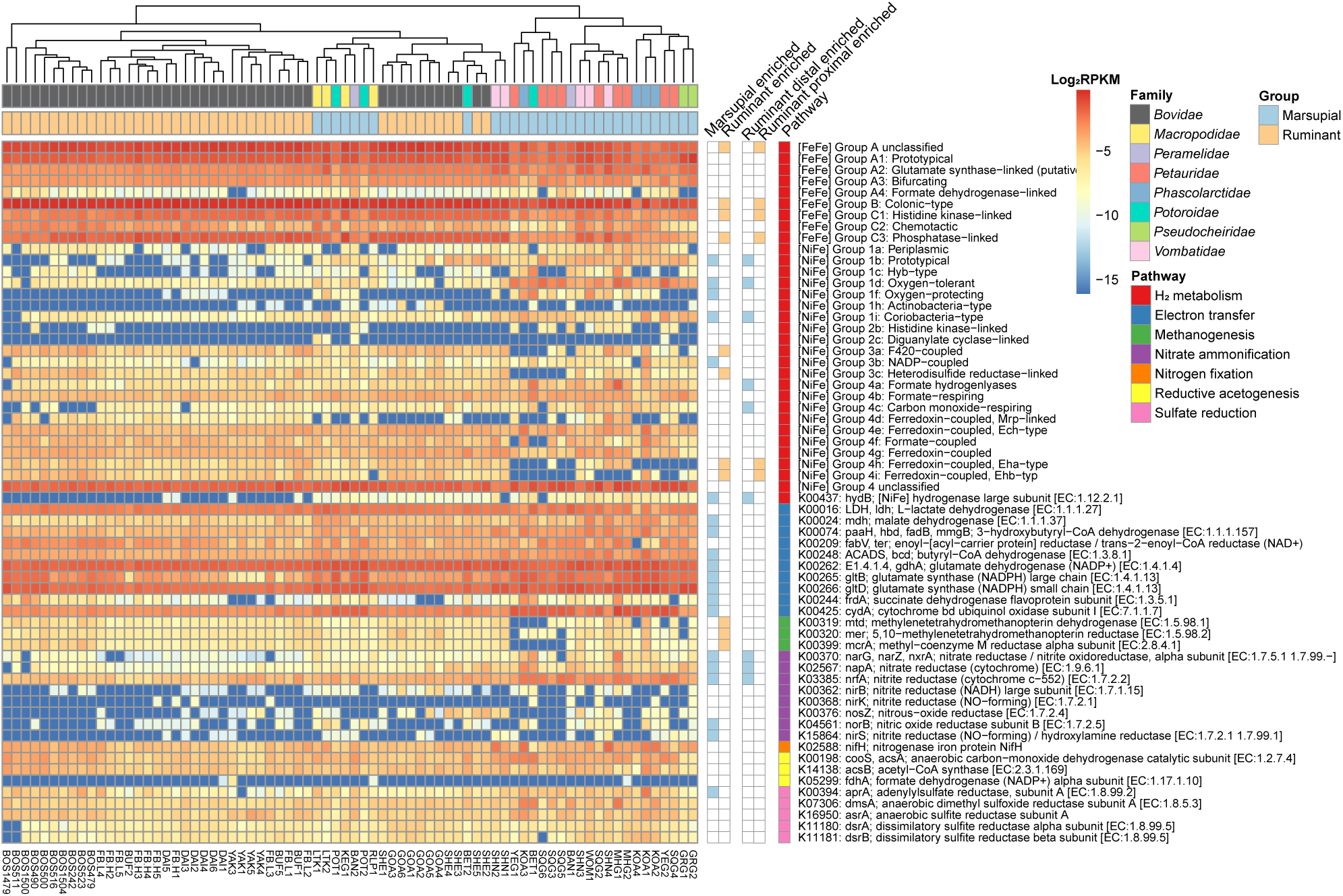
Hydrogen metabolism and reductant disposal gene abundance in marsupial and ruminant faecal samples. Heatmap displays log2-transformed RPKM values based on read mapping to hydrogen cycling proteins from project protein database [32, 39]. Significantly enriched proteins/protein groups are indicated in columns to the right of heatmap.

Although the marsupial and ruminant metagenomes were mostly separated along the primary principal component, some of the marsupial samples more closely resembled ruminants (**Fig. S8A**) and heatmap visualisation confirmed clustering of this subset (n=10) with *Bovidae* samples (**Fig. 3**). Most of these samples were derived from foregut fermenting animals, though the group also includes three hindgut fermenters, namely two Southern hairy-nosed wombats and a Northern brown bandicoot (**Fig. S8A**). Differential abundance analysis revealed enrichment of a subset of the ruminant-linked hydrogenase genes within the ruminant-like marsupials (**Fig. 3, Tables S8 & S9**). We also observed within-marsupial divergence including enrichment of nitrogen cycling proteins glutamate dehydrogenase, glutamate synthase, glutamate synthase-linked hydrogenases and nitrite reductase in the predominantly terrestrial families over arboreal families (**Fig. S8B, Table S10**). The methanogenesis marker *mtd* and F_420_-coupled methanogenesis-linked hydrogenases were also discriminatory along this axis, with enrichment in terrestrial families. Sulfate metabolism enzymes were enriched in the arboreal direction, potentially associated with the enrichment of host glycan-degrading CAZymes in this group and offering an alternative hydrogen sink for their hosts.

### Marsupial methanogens are predicted to use multiple electron and carbon sources

Archaea were estimated to account for an average of <1% of the faecal prokaryotic community based on marker gene-based profiling (**Table S1**, **Fig. 1E**). The highest portion (∼5%) was observed in one Southern hairy-nosed wombat (SHN4, **Fig. 1E**), dominated by *Methanocorpusculum vombati* at 3% of the total community (**Table S11**). Larger archaeal communities were also present in both mahogany gliders (dominant species *Methanocorpusculum petauri*) and one koala (dominant *Methanosphaera* species). Animals with high archaeal abundance typically had communities dominated by either *Methanocorpusculum* spp. or *Methanobacteriaceae* spp. (typically *Methanobrevibacter* spp.) suggestive of competition between these lineages, as previously noted [22]. The most prevalent archaeal species was *Methanocorpusculum petauri*, present in 21% of samples, followed by *Enteromethanobrevibacter sp900769095* (15%), *Methanocorpusculum sp001940805* (12%) and a novel species belonging to the family *Methanomethylophilaceae* (*UBA71* sp., *Candidatus* Methanoprimaticola [40]).

To gain a better understanding of the metabolic capabilities of marsupial methanogens, we reconstructed 38 MAGs supported by targeted long-read sequencing and metabolically annotated them. This revealed genus-level specialisation in substrate preference as previously described [6–9] (**Fig. 4** & **Fig. S9**). The hydrogenotrophic methanogenesis pathway was largely complete in *Methanobrevibacter*, *Methanosphaera* and *Methanocorpusculum* spp., although conservation varied between genomes, particularly for *Methanocorpusculum* representatives (**Fig. S9**). The methanol-based methylotrophic methanogenesis pathway was identified in *Methanosphaera* spp. and *Methanarcanum hacksteinii*, human-derived representatives of the latter previously noted for their lack of complete methanogenesis pathways [40]. The *Ca.* Methanoprimaticola genus MAGs also encoded complete methylotrophic methanogenesis pathways using methanol and di/tri/methylamine, although with non-canonical methyltransferases. There was evidence for ethanol-based methanogenesis in *Methanobrevibacter_B boviskoreani* MAGs [23], with additional alcohol and aldehyde dehydrogenases identified in most genomes. Formate use was also predicted for *Methanocorpusculum* and *Methanobrevibacter* spp. Altogether, these findings suggest that small but taxonomically and metabolically diverse populations of methanogens inhabit marsupials, that use both hydrogen and one-carbon substrates to drive methane production.

**Fig. 4.**
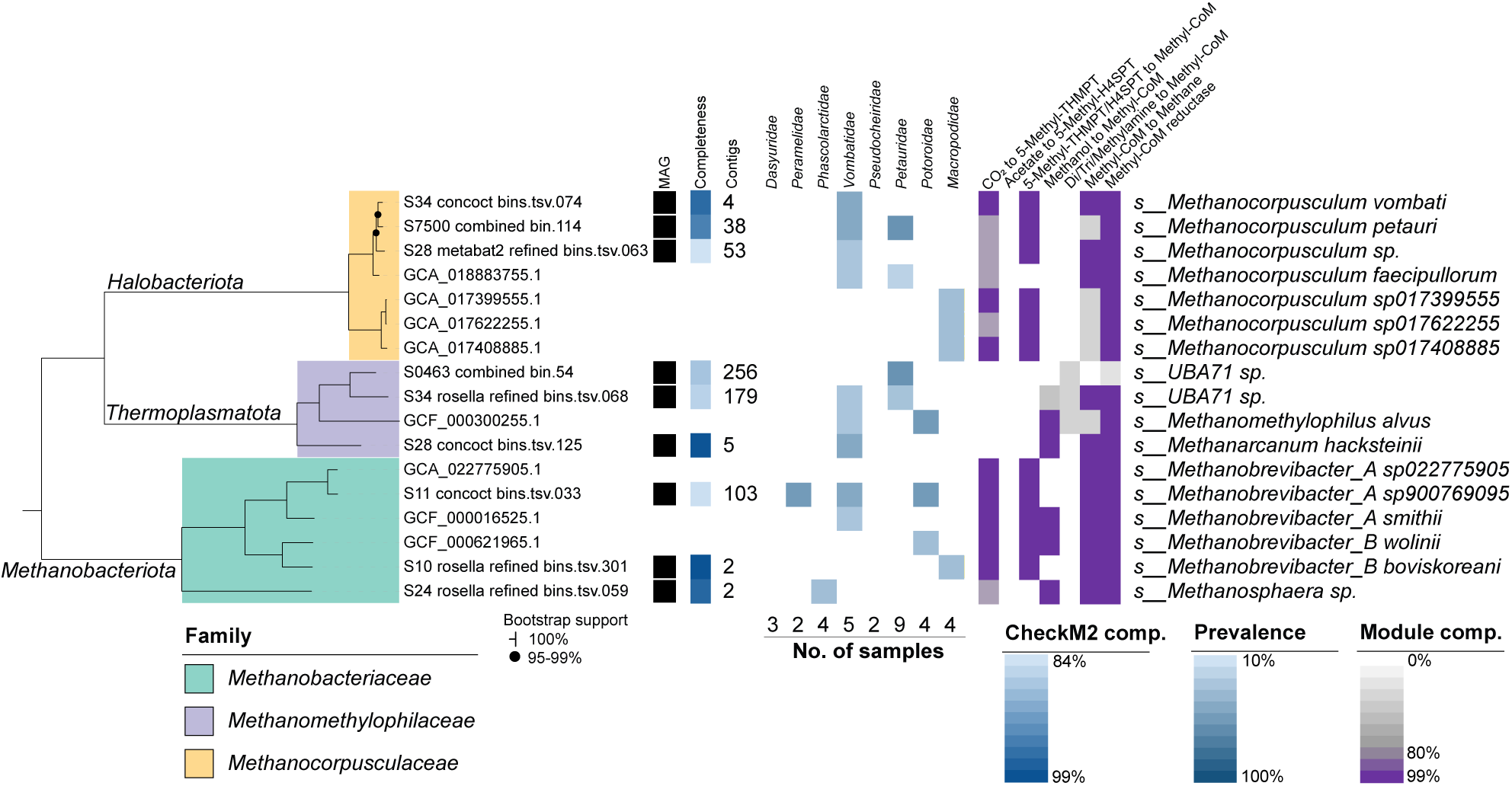
Archaeal genome tree. Maximum likelihood phylogenetic tree inferred from alignment of 53 GTDB single-copy marker genes [70]. Includes genomes from the dereplicated database plus species representative genomes for SingleM-identified species not represented in genome database. Presence in each marsupial host family indicated by coloured boxes and determined based on sample read mapping to genome database with presence cutoffs of >0.05% relative abundance plus >10% genome coverage. Methanogenesis pathway module completeness indicated for the representative genome of each species based on KEGG database annotation.

### Diverse hydrogenotrophic respiratory and acetogenic bacteria inhabit marsupials

Finally, we used the compiled genome database to profile the identities and functions of the hydrogenotrophic bacteria inhabiting marsupials. 33 species encode membrane-bound H_2_-uptake hydrogenases (groups 1a-d, 1f, 1i [NiFe]-hydrogenases) that input electrons into the respiratory chain (**Table S12**). As expected, these species include *Desulfobacterota* (e.g. *Bilophila*, *Desulfovibrio*, group 1a) encoding the pathway for dissimilatory sulfate reduction (i.e. Apr and Dsr enzymes), with hydrogenotrophic nitrate-reducing *Campylobacter_D* also detected (group 1b). Also present were 11 species predicted to mediate hydrogenotrophic dissimilatory nitrate reduction to ammonium (DNRA, *via* Nar and Nrf enzymes, [41]) including *Actinomycetota* species *Eggerthella lenta* and *Raoultibacter massiliensis*, and *Desulfobacterota* species *Bilophila wadsworthia* and *Desulfovibrio piger*. Other inferred hydrogenotrophs include *Parabacteroides*, *Megamonas*, *Parasutterella*, *Akkermansia*, and the uncultivated *Planctomycetota* genus *RUG369*. Culture-based experiments are needed to confirm the role of hydrogen uptake in each of these genera and whether their introduction or stimulation in the rumen could help to reduce methane levels. Consistent with these findings, four of the five hydrogenase subgroups (groups 1b, 1d, 1f, 1i) encoded by these taxa and DNRA genes were statistically enriched in marsupials compared to ruminants (**Fig. 3, Table S8)**. Together, these diverse bacteria are likely to coordinate and govern metabolic schemes that can, with the availability of inorganic electron acceptors, feasibly reduce gut hydrogen levels and thereby promote a low methane economy.

Given the known association of acetogens with macropods [34, 35] and stimulation of these lineages following methanogenesis inhibition in ruminants [16], we examined acetogenic populations across marsupials. Seventy species from 23 genera carried both the *acsB* and *cooS* genes encoding the key determinants for acetogenesis, the acetyl-CoA synthase/carbon monoxide dehydrogenase complex (**Table S12**). Most marsupial hosts harboured multiple acetogenic genera except for the greater glider, which were underrepresented in the genome database, and the macropods, carrying solely *Ca.* Faecousia species. All predicted acetogens affiliated with five families of *Clostridia*, except for the putative flagellate-associated acetogen *Candidatus* Adiutrix from the phylum *Desulfobacterota* [42]. MAGs were obtained for the well-characterised acetogen genus *Blautia* as well as numerous uncultivated lineages, most notably multiple MAGs of *Ca.* Faecousia and *CAG-170* that we previously showed become enriched in ruminants following methanogenesis inhibition [16]. Almost all (90%) of these species encode [FeFe]-hydrogenases from group A (29 species) and/or B (57 species), suggesting that they can use H_2_ to reduce CO_2_ to acetate. It is unclear how acetogens co-exist and potentially outcompete methanogens in marsupials given their higher H_2_ threshold and lower energy yield [43]. One possibility is that these acetogens adopt a mixotrophic metabolism, using a combination of organic and inorganic substrates, including direct formate assimilation rather than using the hydrogen-dependent CO_2_ reductase reaction as recently described [16]. It is also possible that the spatial organisation of marsupial guts provides a range of niches that enable co-existence of acetogens and methanogens.

## Discussion

The rationale and justification that marsupials are modest methane emitters compared to ruminants is based on a small number of studies detailing low methane emissions relative to digestible feed intake and/or a low abundance of methanogenic archaea and bacteria offering alternative hydrogen sinks [17–22, 25]. However, broad comparison of methanogen populations across marsupial hosts, particularly beyond members of the *Macropodidae*, is lacking. Here, we provide a metagenomic analysis of the faecal communities across 14 marsupial species representing eight families, with a focus on methanogens and metabolic pathways that provide alternative electron sinks to methanogenesis. Two types of methanogens persist: hydrogenotrophic (genera *Methanobrevibacter*, *Methanosphaera* and *Methanocorpusculum*) and methylotrophic (genera *Methanosphaera*, *Methanarcanum,* and *Ca.* Methanoprimaticola). The most prevalent archaeal species was *Methanocorpusculum petauri*, present in 21% of samples, the majority of these originating from namesake *Petauridae* hosts [22]. Most methanogenic archaea were identified in one or two host families, with only *Enteromethanobrevibacter sp900769095* found in three, suggestive of host specificity, as previously observed in gut archaea [44]. *Vombatidae* hosts carry the highest diversity of methanogens, followed by members of family *Petauridae* and *Macropodidae*. However, we note that the diversity of identified species per host in this study differs from previous analyses. For example, we did not identify *Methanobrevibacter* or *Methanocorpusculum* species in koala, in contrast to previous studies [21, 22]. The limited sample size and observed high variability between individual animals is expected to contribute to such differences.

The archaeal community did not differ significantly across marsupial hosts, with the latter likely being a product of both the low relative abundance and high variability of archaea between animals, as seen in marsupial faecal fungal communities [45]. In contrast to the archaeal community, the bacterial community divergence was influenced by the confounded variables of host species, family, gut type (foregut/hindgut/simple hindgut) and diet (omnivore/herbivore/carnivore), with host species/family explaining the majority of variation. We observed clustering of predominantly grazing animals from different families (*Vombatidae*, *Macropodidae*, *Potoroidae*) in a comparison of all hosts, driven by the relative abundance of members of the genus *Ca.* Faecousia, suggesting a common diet is influencing community structure. *Ca.* Faecousia is an as-yet-uncultured genus belonging to the *Bacillota* family *Oscillospiraceae* commonly found in faecal samples and named as part of a large MAG-based study of chicken faeces [36, 46]. Members of this predicted acetogenic genus, along with acetogenesis pathway markers, are enriched in dairy calves in response to feeding with the methanogenesis inhibitor 3-nitrooxypropanol (3-NOP), indicating a shift toward H_2_ disposal via acetogenesis with this treatment [16]. A similar increase in the acetogenesis markers was observed in related studies, though in association with different taxa, indicating the functional effect as dominant and the driving taxa as dependent on the specific gut community [16]. Acetate production has been confirmed as a primary output of macropod foregut communities [34, 47] and is considered the dominant H_2_ disposal mechanism in these animals [48]. While we identified multiple acetogenic genera in most marsupial hosts, only *Ca.* Faecousia species were confirmed in macropodid hosts. Proposed association of acetogens with epithelial-attached biofilms in the macropodid forestomach [48] may limit their recovery via faecal sampling.

We did not see any significant differences in reductive acetogenesis markers between terrestrial and arboreal marsupials, supporting similar acetogenic capacity across their different diets. The terrestrial animals, however, were enriched in glutamate and nitrogen metabolism markers, in combination with methanogenesis markers, suggesting these pathways are competing for electrons with H_2_ formation in some marsupial hosts and pointing to potentially increased methanogenesis in terrestrial vs arboreal marsupials. While methane production has been quantified in wallabies and kangaroos [20], methane production kinetics and intensity is unknown for other marsupials. However, anaerobic culturing of faecal samples from multiple marsupial hosts supports increased methanogenic capacity of terrestrial over arboreal animal gut communities [22].

Despite most marsupials harbouring multiple acetogenic species, we found no difference in the abundance of acetogenesis markers between marsupials and ruminants. Marsupial samples, however, were enriched in other alternative hydrogen uptake and disposal pathways, including butyrate, propionate and glutamate metabolism. Marsupial hosts were also enriched in nitrate and nitrite reductases supporting increased nitrate ammonification in these hosts. Enrichment of these alternative hydrogen sinks in marsupials supports reduced availability of substrates for methanogenesis in these animals [39, 49–51]. These findings mirror those comparing the rumen microbiome of low- to high-methane-emitting cattle, which identified enrichment of butyrate, propionate, glutamate and nitrate ammonification genes in low-methane-emitting cattle [39]. Conversely, methanogenesis genes and methanogenic hydrogenases were enriched in high-methane-emitting cattle [39]. We also observed enrichment of cytochrome *bd* oxidase and oxygen-tolerant hydrogenases in marsupials relative to ruminants, supporting the presence of low levels of oxygen in the marsupial gut that may inhibit methanogens and the use of alternative electron sinks.

The marsupial faecal community also displays enrichment in pectin degradation enzymes in comparison to ruminants. Bacterial pectin degradation can produce methanol [53], providing a substrate for methylotrophic methanogens. The methanol-based methylotrophic methanogenesis pathway was identified in *Methanosphaeara* spp. and *Methanarcanum hacksteinii,* which are present in some hindgut marsupials and could potentially be supported by pectin degradation. While pectin degradation occurs in ruminants, methane production from methanol increases following exposure to a pectin-containing diet, suggestive of microbial adaptation and therefore diet influencing the degree of pectin utilisation [54]. Starch degrading enzymes were enriched in ruminants relative to marsupials, which may be associated with agricultural ruminant diets high in starch that is not fully degraded in the rumen [55].

While these data support a low methane-emitter prediction for marsupials based on the functional profile of their gut microbiomes, we acknowledge this conclusion is limited by the low number of animals included in the study, their predominantly captive status and a lack of *in vivo* metabolite measurements (e.g. hydrogen and methane). Known differences in gut microbiome composition exist between wild and captive animals [56], therefore these conclusions need to be validated in wild animals. We are also limited to considering faecal samples, which likely do not reflect the complete functional profile of the gut community, particularly regarding foregut-fermenting species. While we only included faecal and rectal samples in comparing marsupials to true ruminant hosts, analysis of marsupial foregut samples is necessary to complete the picture. Despite these limitations, the consistent parallels between the functional profiles of marsupial and ruminant faecal communities versus that of low- and high methane-emitting ruminants supports the predicted low-methane phenotype of marsupials and suggests that they could be used as natural models of low methane emitting gut microbiomes.

## Methods

### Sample collection

Faecal samples were collected from 14 species of marsupial from predominantly captive environments in Queensland, Australia (**Table S1**). Species included were the fat-tailed dunnart (*Sminthopsis crassicaudata*, n=3), Lumholtz’s tree kangaroo (*Dendrolagus lumholtzi*, n=2), Eastern grey kangaroo (*Macropus giganteus*, n=1), red-legged pademelon (*Thylogale stigmatica*, n=1), Northern brown bandicoot (*Isoodon macrourus*, n=2), yellow glider (*Petaurus australis*, n=2), mahogany glider (*Petaurus gracilis*, n=2), squirrel glider (*Petaurus norfolcensis*, n=5), koala (*Phascolarctos cinereus*, n=4), rufous bettong (*Aepyprymnus rufescens*, n=2), long-nosed potaroo (*Potorous tridactylus*, n=2), greater glider (*Petauroides volans*, n=2), Southern hairy-nosed wombat (*Lasiorhinus latifrons*, n=4) and common wombat (*Vombatus ursinus*, n=1). Faecal samples were stored at −80°C as soon as possible following collection. Ethical permission for the collection of all samples was granted by the Animal Ethics Unit, The University of Queensland, Brisbane, Australia, under ANRFA/SCMB/099/14 and ANRFA/SCMB/475/20. Public ruminant datasets incorporated into this study are available under NCBI BioProjects PRJNA657455 [57], PRJNA624740 [58], PRJNA752224 [59] and PRJNA987743 [60]. Only faecal or rectal samples were included (**Table S13**).

### Sample preparation and metagenomic sequencing

Total genomic DNA was extracted from approximately 60-200 mg of faecal material. Samples were homogenised at 2,000 rpm for 5 min using the MoBio PowerLyzer24 in a MoBio bead tube containing 0.1 mm diameter zirconia/silica beads and 750 µL of TLA buffer (Promega, WI, USA). Supernatant was recovered via centrifugation at 10,000 g for 30 s. DNA was extracted from 150 µL of supernatant using the Maxwell 16 robotic system and corresponding Tissue DNA kit (Promega, WI, USA) following the manufacturer’s instructions.

Illumina sequencing libraries were prepared using the Nextera DNA Library Preparation kit (Illumina, CA, USA). Samples were sequenced on the NovaSeq6000 platform, with some samples supplemented with data generated using the HiSeq2000 platform, producing ∼ 7 Gb of 150 bp paired-end reads per sample.

Oxford Nanopore sequencing was performed on a PromethION instrument using an R10.4.1 flow cell following sample preparation with the Ligation Sequencing Kit SQK-NBD114-24 (Oxford Nanopore, UK). Samples were multiplexed using native barcoding and run using adaptive sampling generating ∼5 Gb per sample with an average read length of 800 bp. Five archaeal MAGs were used as a positive filter for adaptive sampling using MinKNOW v23.07.5. Enrichment of archaeal populations was estimated by comparison of their relative abundance between Illumina and Nanopore sequencing runs (**Table S14**). Guppy v7.0.9 was used for initial base-calling in high accuracy mode. Subsequent recalling to super high accuracy was undertaken using Dorado v0.5.2 (https://github.com/nanoporetech/dorado) following pod5 conversion with pod5 v0.3.10 (https://github.com/nanoporetech/pod5-file-format).

### Data assembly

Illumina data was quality trimmed and cleaned of adapters using Trimmomatic v0.39 [61]. Read counts pre- and post-trimming for all data were generated using SeqKit v2.4.0 [62]. The trimmed reads were then assembled with MEGAHIT v1.2.9 [63]. Reads were mapped to assembled contigs using BamM (https://github.com/Ecogenomics/BamM) and binned using MetaBAT2 v2.15 [64].

Sequence barcodes were removed from Nanopore sequence data using Porechop v0.2.4 (https://github.com/rrwick/Porechop), reads with mid-strand barcodes were also removed. Hybrid assembly of Nanopore and Illumina data from the same samples was undertaken using Aviary v0.8.3 (https://github.com/rhysnewell/aviary) with default parameters. Binning was performed by Aviary v0.8.3 employing Rosella v0.5.3 (https://github.com/rhysnewell/rosella), MetaBAT1 and MetaBAT2 v2.15 [64] and CONCOCT v1.1.0 [65] based on read mapping of all five Nanopore-targeted samples to each assembly. DAS Tool v1.1.6 [66] was used to determine a non-redundant MAG set.

All MAGs were quality assessed using CheckM v1.2.2 [67] and CheckM2 v1.0.2 [68] and classified using GTDB-Tk v2.3.0 [69] against GTDB releases 08-RS214 and 09-RS220 [70, 71]. Nanopore- and Illumina-based MAGs exceeding 50% completeness with ≤5% contamination according to either CheckM or CheckM2 were combined and dereplicated at 99% average nucleotide identity (80% alignment fraction) using CoverM v0.7.0 (https://github.com/wwood/CoverM) applying the ‘Parks2020_reduced’ quality formula.

### Genome database

Public genomes to be included in the final genome database were selected from those within GTDB release 08-RS214 [70, 71] based on relative abundance and coverage values generated using CoverM v0.7.0 (https://github.com/wwood/CoverM). Genomes were included that achieved relative abundance of >0.05% and >0.1X coverage in at least one sample. Selected genomes were combined with the dereplicated sample MAGs (≥50% complete with ≤5% contamination) and the combined set dereplicated at 95% average nucleotide identity (60% alignment fraction) using CoverM v0.7.0 applying the ‘Parks2020_reduced’ quality formula. Community profiling was undertaken using relative abundance values generated from mapping short-read data to this database using CoverM v0.7.0 ‘genome’. BWA [72] implemented within CoverM was used for Illumina read mapping with the parameter ‘-k 31’, while Minimap2 [73] option ‘-p minimap2-hifi’ was used for Nanopore data (used for host DNA estimation and evaluation of adaptive sequencing only). Read alignments were filtered for those achieving ≥95% identity across ≥90% of the read length. Genomes were functionally annotated using DRAM v1.5.0 [74] using the KOfam [75], Pfam [76] and dbCAN [77] databases accessed 21 February 2024. Contigs not recruited into MAGs were compiled for assessment of the taxonomic composition of unmapped reads, using MMseqs2 v15-6f452 [78] ‘easy-taxonomy’ against the NCBI NR protein database [79] for taxonomic inference.

### Host tree

The marsupial host phylogenetic tree was inferred using three nuclear (*IRBP, BRCA1, vWF*) and two mitochondrial genes (*CYTB, ND2*), which were selected based on sequence availability (**Table S15**). Sequences were aligned using MAFFT v7.490 [80] and trimmed of gaps present in >10% of sequences using trimAl v1.4.1 [81]. Maximum likelihood trees were inferred using IQ-TREE v2.2.2.3 [82] using ModelFinder for model selection with gene and codon partitioning. Bootstrap support was generated from 100 replicates.

### Host DNA estimation

An estimate of host DNA contamination was created using available marsupial genomes in combination with a set of exon capture sequences [83] (**Table S16**). Due to the absence of full-length genomes for all hosts, this analysis is considered to provide only a minimum estimate of host DNA contamination. Read mapping was undertaken using BWA [72] as above. Estimated host DNA was calculated based on read numbers aligning with ≥95% identity across ≥90% of the read length determined using CoverM v0.7.0.

### Marker gene-based community profiling

SingleM v0.16.0 [36] was used for marker gene-based community profiling using the GTDB release 08-RS214 metapackage, with the ‘condense’ function used to summarise data across markers and SingleM microbial_fraction used to estimate the bacterial and archaeal community fraction [84].

### Protein database creation and functional annotation

The gene database was created from proteins within assembled contigs from marsupial and ruminant samples annotated using Prodigal v2.6.3 [85]. Proteins were clustered at 90% identity across 80% of protein length using MMseqs2 v14-7e284 [78]. Representative protein sequences were functionally annotated using DRAM v1.5.0 [74] using the KOfam [75], Pfam [76] and dbCAN [77] databases accessed 21 February 2024.

Protein abundance within samples was calculated based on read alignment using DIAMOND v2.1.0 [86] to the protein database, filtering for sample reads ≥ 140 bp with Seqkit v2.4.0 [62] (excepting public dataset PRJNA987743 where read length was 120 bp). Alignments were filtered using settings evalue 0.00001, min-score 40, query-cover 80, id 70, max-hsps 1 and max-target-seqs 1. Alignment of reads to the single copy marker genes included within the SingleM GTDB release 08-RS214 metapackage [36] was undertaken using the same settings. Reads per kilobase per million mapped read (RPKM) values were calculated for both the protein database and SingleM marker genes, with the mean RPKM value across all SingleM markers used to normalise protein database values on a per sample basis.Normalised RPKM values per protein were summed across equivalent functional annotations (e.g. KEGG orthologs or hydrogenase groups) and log2 transformed for PCA and differential abundance analysis.

### Hydrogenase classification

Pfam annotations from DRAM were used to collate a set of hydrogenase proteins present in the protein database or in MAGs based around hydrogenases listed in Greening, Geier [32] (**Table S17**). Collated protein sequences from the protein database were clustered at 50% identity and representatives submitted to HydDB for classification [87]. Taxonomic origin of hydrogenases (and KOs of interest) was inferred using MMseqs2 v15-6f452 [78] ‘easy-taxonomy’ on containing contigs (database: GTDB 08-RS214).

### Methanogenesis pathway analysis

Archaeal genomes were annotated using DRAM v1.5.0 [74]. KEGG module presence was determined using EnrichM v0.6.1 (https://github.com/geronimp/enrichM) with custom modules describing sub-components of methanogenesis pathways (**Table S18**). Candidate missing enzymes were identified using BLAST [88] with characterised reference sequences from UniProt [89].

### Statistical analyses and figure generation

PCA was conducted using vegan v2.6-4 (R v4.3.2) [90] using Hellinger-transformed data following filtering for taxa present in ≥2 samples. Contributing variables were filtered and plotted using factoextra v1.0.7 (R v4.3.2) [91]. All other R-based analyses were conducted using R v4.4.0. Redundancy analysis was conducted using the ‘rda’ function within vegan v2.6-6.1, with significance determined using ‘anova.cca’ [90].

Differential abundance analysis was undertaken using three methods: sPLS-DA implemented within mixOmics v6.28.0 [92] and linear regression-based methods implemented within MaAsLin2 v1.18.0 [93] and LinDA/MicrobiomeStat v1.2 [94]. Data was filtered for taxa present in ≥2 samples. Relative abundance values were transformed via centred-log transformation for sPLS-DA and regression analyses. sPLS-DA was run using 5-fold cross validation with 50 repeats, using the ‘tune’ function to select the number of taxa to include. MaAsLin2 and LinDA were run with gut type as a fixed effect and species as a random effect.

Boxplots and barcharts were created using ggplot2 v3.5.1 [95]. Heatmaps were created using pheatmap v1.1.12 [96]. Phylogenetic tree figures were created using iTOL [97].

## Data availability statement

Sequencing data is available via NCBI under BioProject PRJNA1167890. Additional sequencing data was produced for three previously published samples: S29, S30 and S34 [22] (**Table S1**). MAGs over 90% complete are available via NCBI with the complete set available at figshare 10.6084/m9.figshare.27168006.v1.

## Acknowledgements

The authors thank Cairns Tropical Zoo, Currumbin Wildlife Sanctuary, David Fleay Wildlife Park, Hidden Vale Research Station, Lone Pine Koala Sanctuary, Wildlife Habitat Port Douglas, A. Shima and D.S. Teakle for the provision of marsupial faecal samples.

## Ethics

Approval for the collection of all samples was granted by the Animal Welfare Unit, the University of Queensland, Brisbane, Australia, under ANRFA/SCMB/099/14.

---

**Fig. S1.**
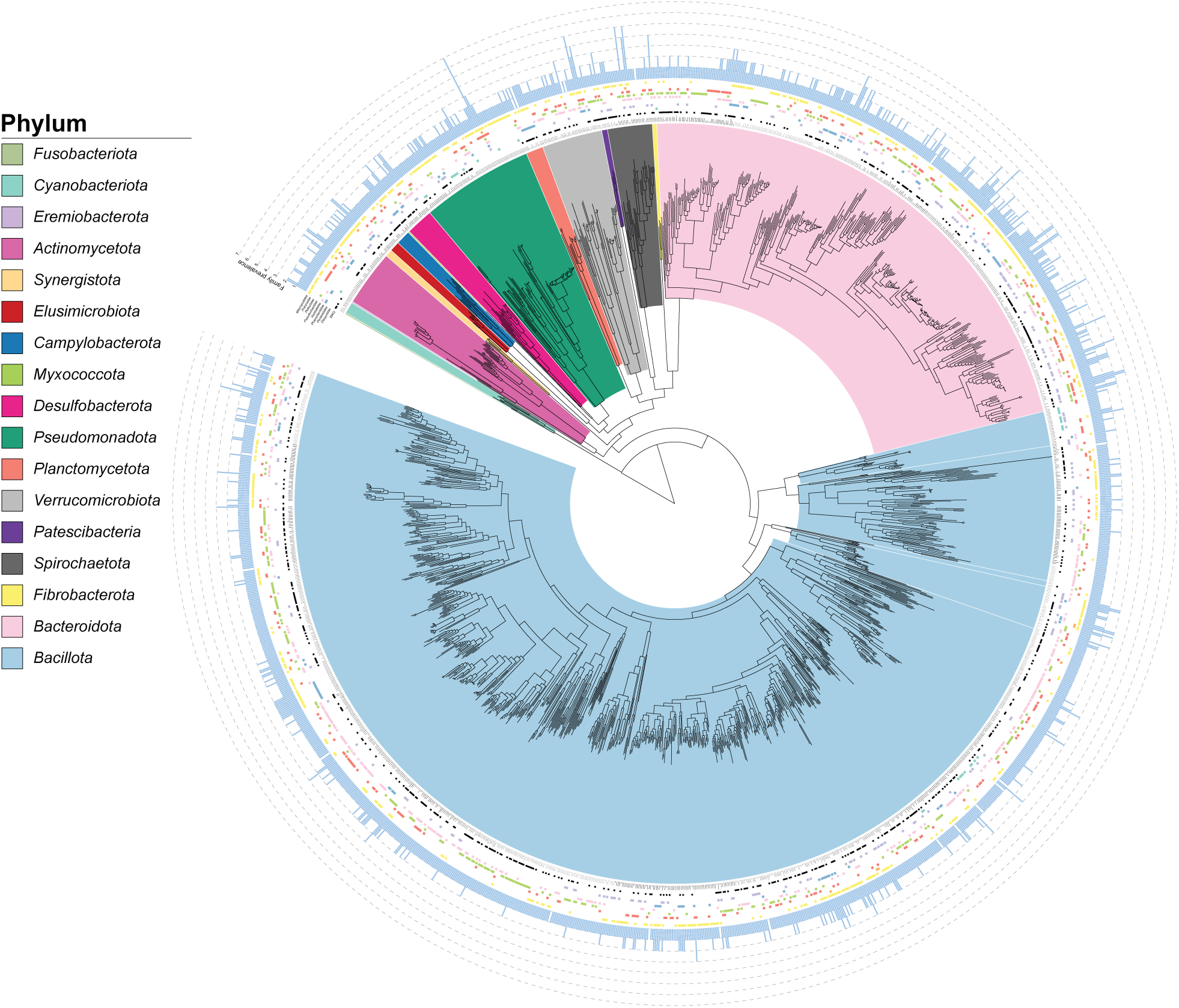
Bacterial genome tree. Maximum likelihood phylogenetic tree inferred from alignment of 120 GTDB single-copy marker genes [69]. Includes genomes from the dereplicated database plus species representative genomes for SingleM-identified species not represented in genome database. Outer rings represent presence in each marsupial host family determined based on sample read mapping to genome database with presence cutoffs of >0.05% relative abundance plus >10% genome coverage. Outermost ring displays count of host families where at least one sample met this threshold.

**Fig. S2.**
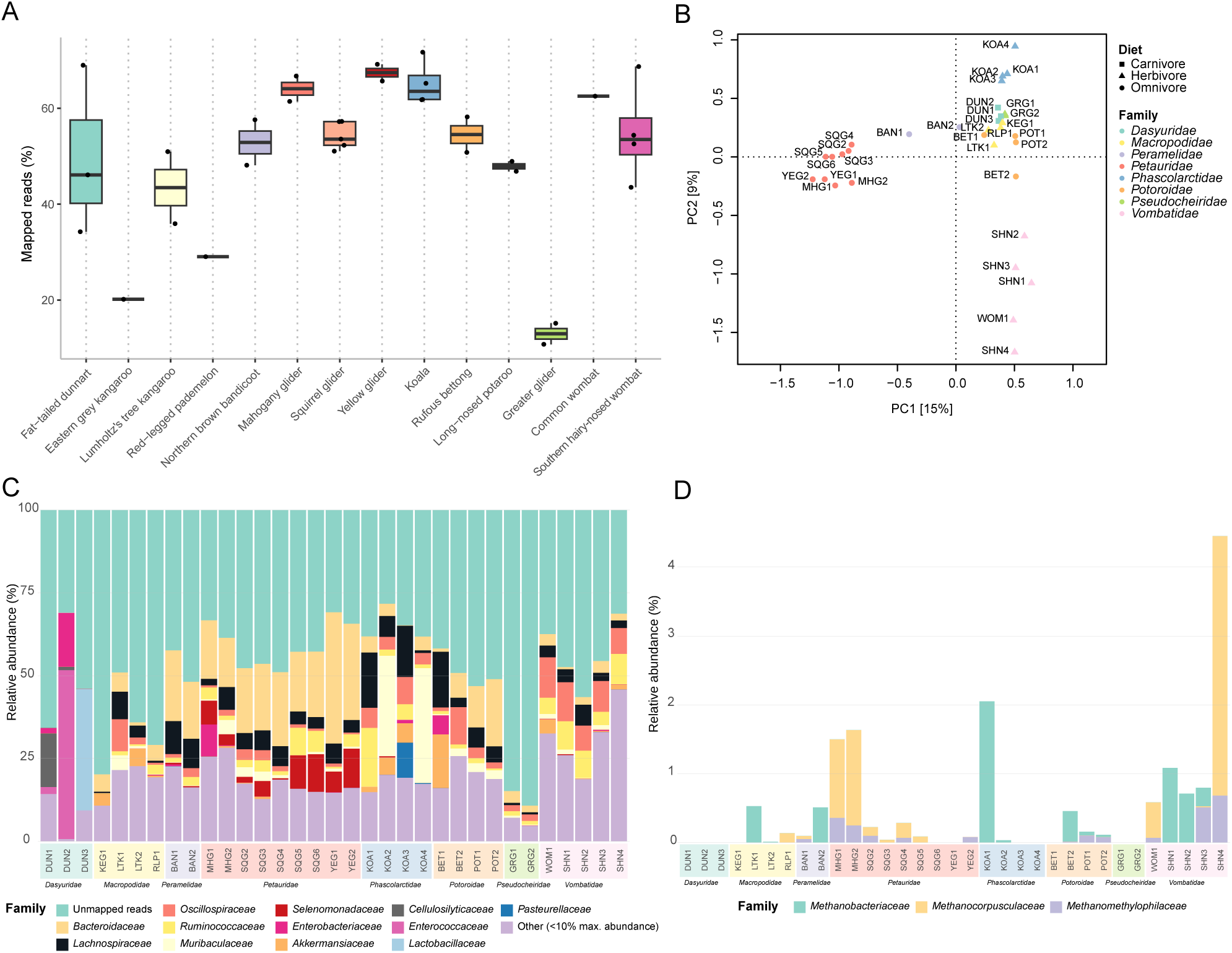
Genome-based community composition. (A) Read mapping to the dereplicated genome database, filtering for alignments >95% identical across >90% of the read length. (B) Principal component analysis based on square-root transformed relative abundance values. Family level relative abundance of bacterial (C) and archaeal communities (D). Host species are abbreviated as DUN: fat-tailed dunnart, KEG: eastern grey kangaroo, LTK: Lumholtz’s tree kangaroo, RLP: red-legged pademelon, BAN: northern brown bandicoot, MHG: mahogany glider, SQG: squirrel glider, YEG: yellow glider, KOA: koala, BET: rufous bettong, GRG: greater glider, WOM: common wombat, SHN: southern hairy-nosed wombat.

**Fig. S3.**
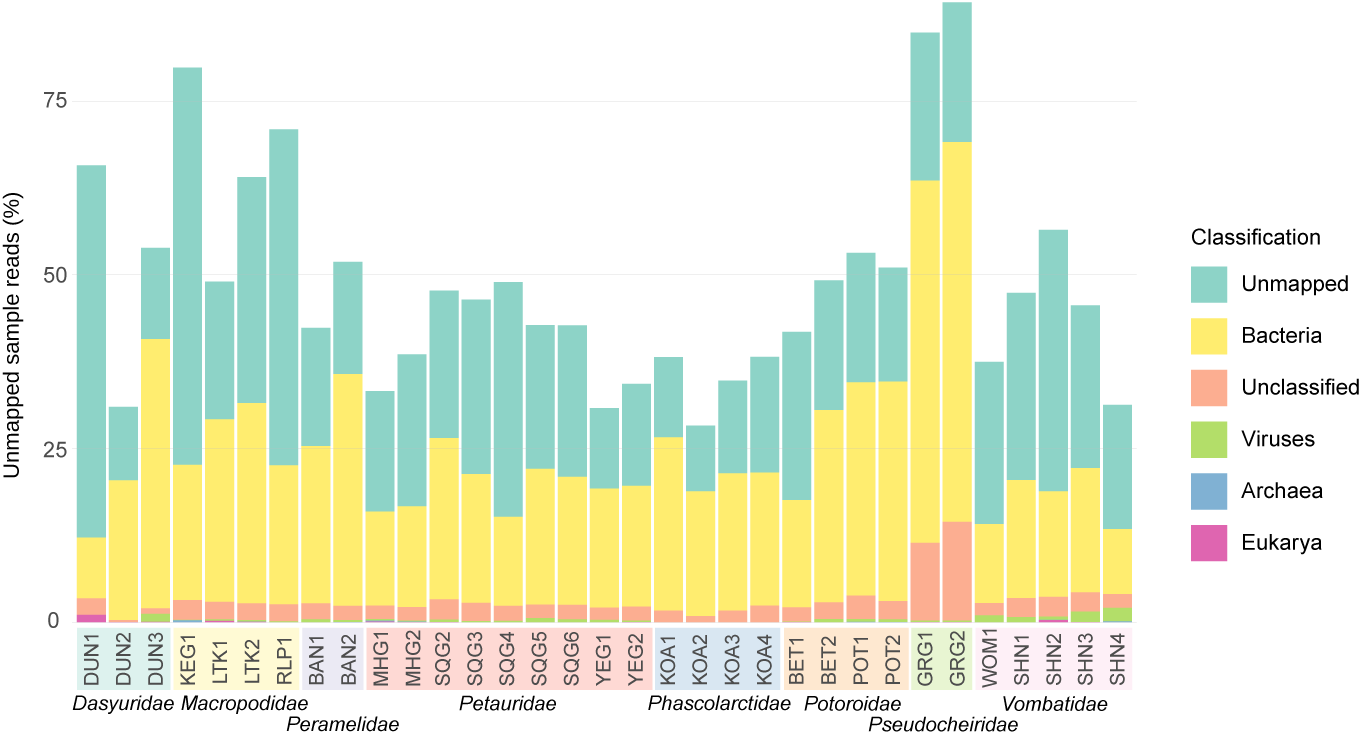
Classification of unmapped sample reads. Sample reads not aligning to the genome database were mapped to assembled contigs not included within study MAGs. Contigs were classified to domain level using MMseqs2 [78] to estimate fractional representation of each domain within the unmapped data. Classification ‘Unmapped’ denotes reads that did not align to the unbinned contigs. Host species are abbreviated as DUN: fat-tailed dunnart, KEG: eastern grey kangaroo, LTK: Lumholtz’s tree kangaroo, RLP: red-legged pademelon, BAN: northern brown bandicoot, MHG: mahogany glider, SQG: squirrel glider, YEG: yellow glider, KOA: koala, BET: rufous bettong, GRG: greater glider, WOM: common wombat, SHN: southern hairy-nosed wombat.

**Fig. S4.**
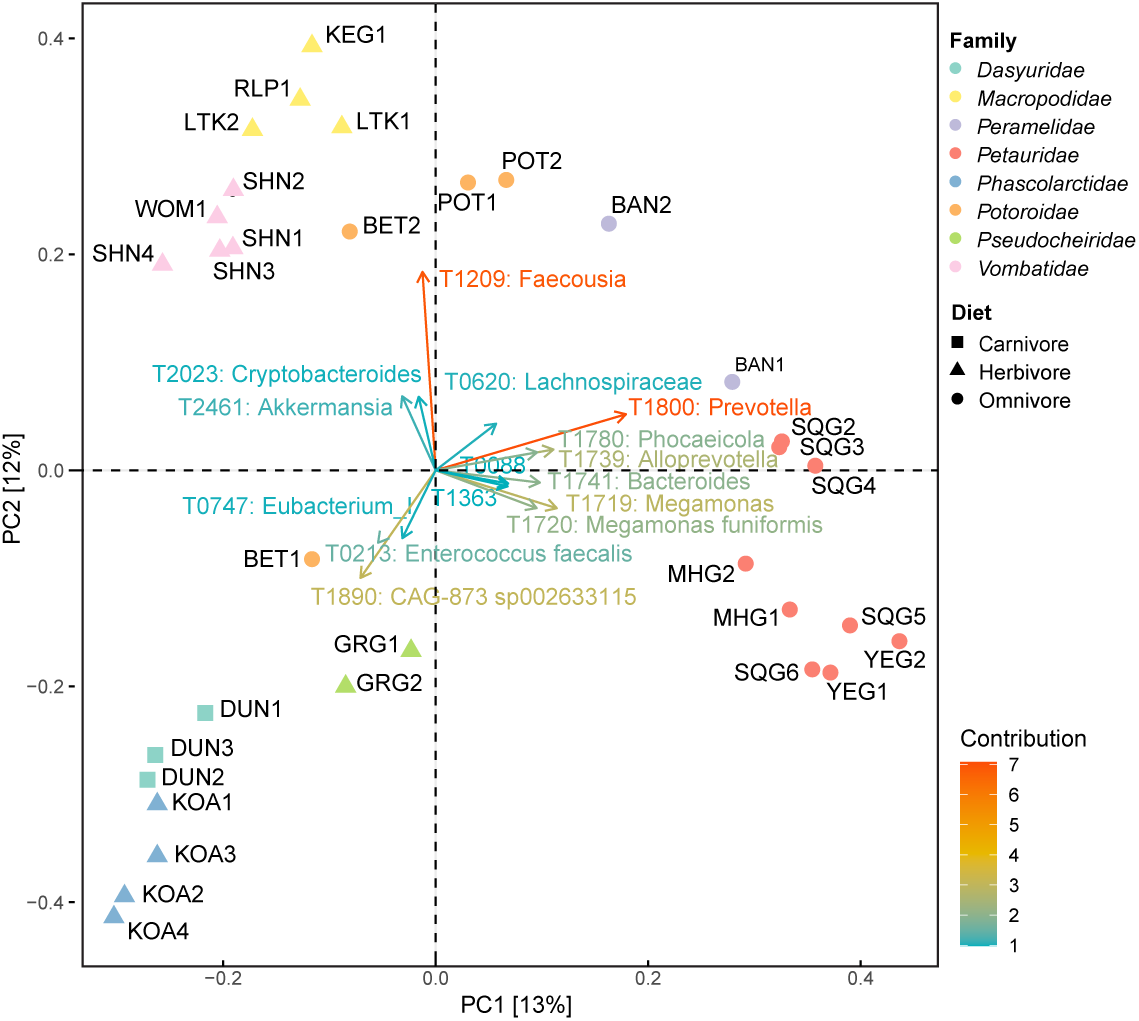
Taxa contributing to marker gene-based community profile components. Principal components analysis of faecal bacterial and archaeal community displaying the top 15 taxa contributing to sample separation. Host species are abbreviated as DUN: fat-tailed dunnart, KEG: eastern grey kangaroo, LTK: Lumholtz’s tree kangaroo, RLP: red-legged pademelon, BAN: northern brown bandicoot, MHG: mahogany glider, SQG: squirrel glider, YEG: yellow glider, KOA: koala, BET: rufous bettong, GRG: greater glider,

**Fig. S5.**
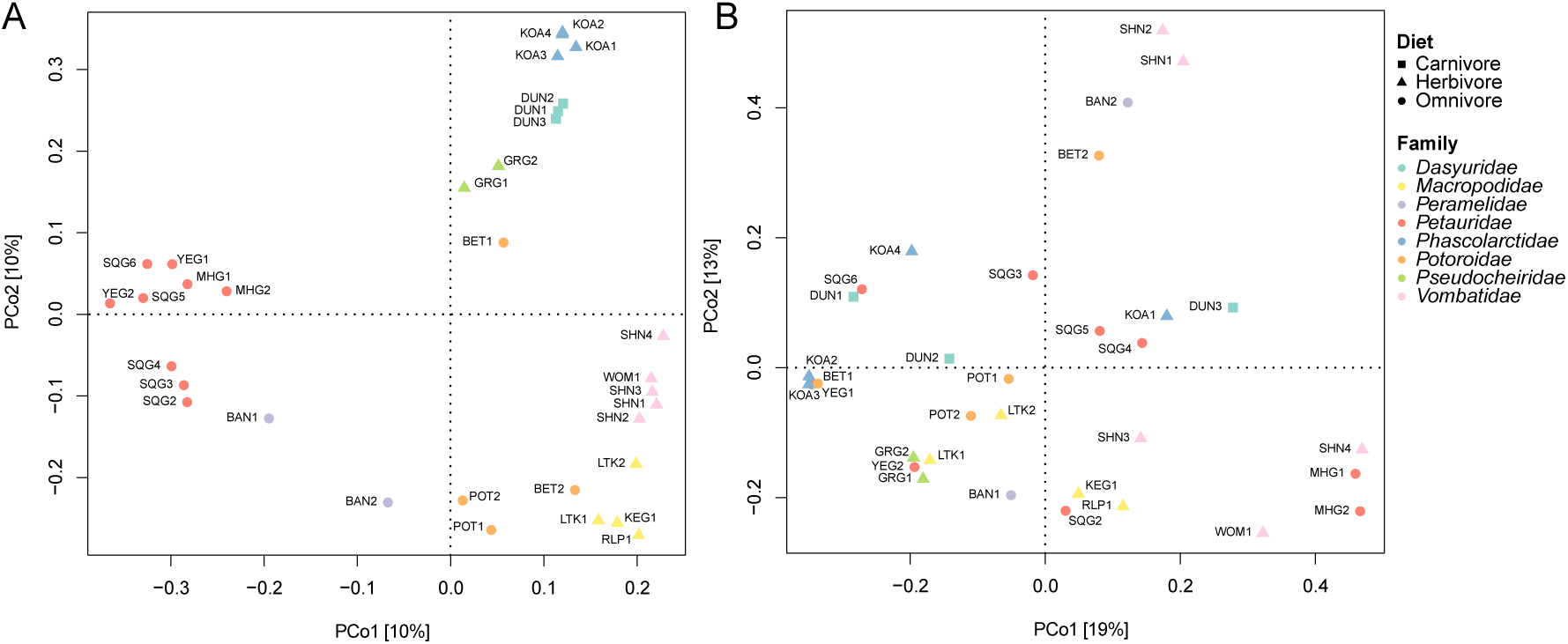
Marker gene-based community presence/absence profiles. Principal coordinates analysis of faecal bacterial (A) and archaeal (B) community presence/absence (Jaccard distance). Host species are abbreviated as DUN: fat-tailed dunnart, KEG: eastern grey kangaroo, LTK: Lumholtz’s tree kangaroo, RLP: red-legged pademelon, BAN: northern brown bandicoot, MHG: mahogany glider, SQG: squirrel glider, YEG: yellow glider, KOA: koala, BET: rufous bettong, GRG: greater glider, WOM: common wombat, SHN:

**Fig. S6.**
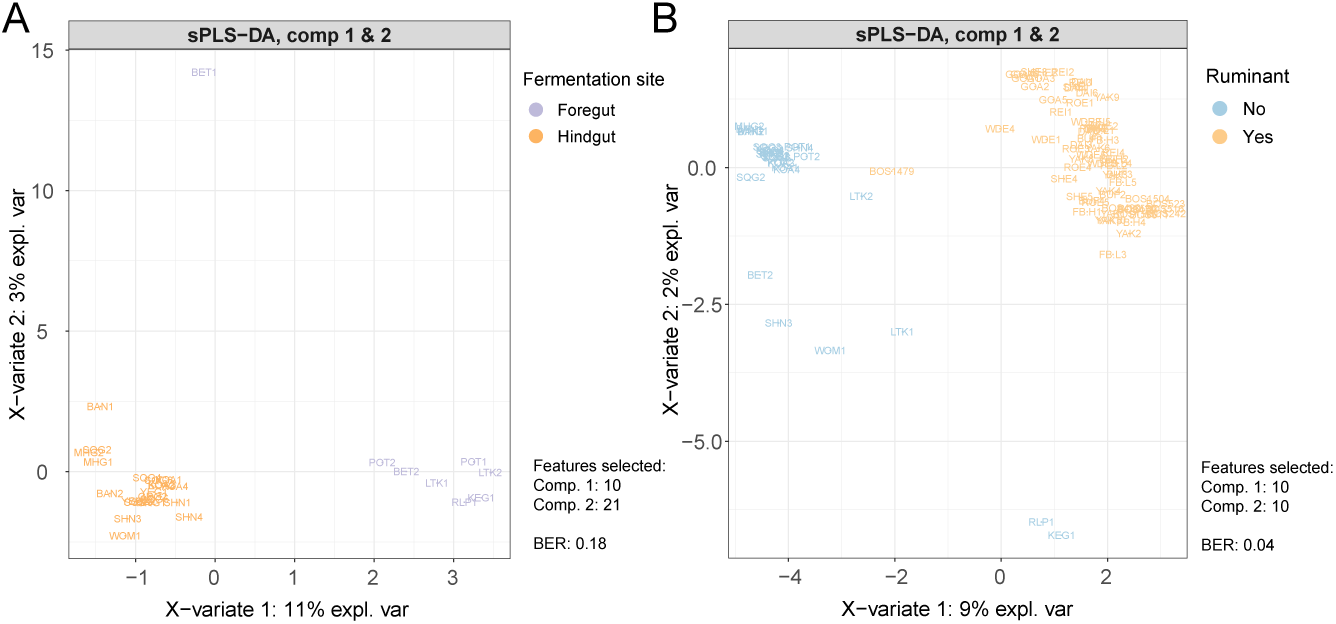
sPLS-DA based on SingleM community profiles. Mixomics-implemented sPLS-DA was carried out on foregut vs hindgut-fermenting marsupials (A), all marsupials vs publicly available faecal/rectal ruminant samples (B) and foregut-fermenting marsupials vs faecal/rectal ruminant samples (C). Host species are abbreviated as DUN: fat-tailed dunnart, KEG: eastern grey kangaroo, LTK: Lumholtz’s tree kangaroo, RLP: red-legged pademelon, BAN: northern brown bandicoot, MHG: mahogany glider, SQG: squirrel glider, YEG: yellow glider, KOA: koala, BET: rufous bettong, GRG: greater glider, WOM: common wombat, SHN: southern hairy-nosed wombat.

**Fig. S7.**
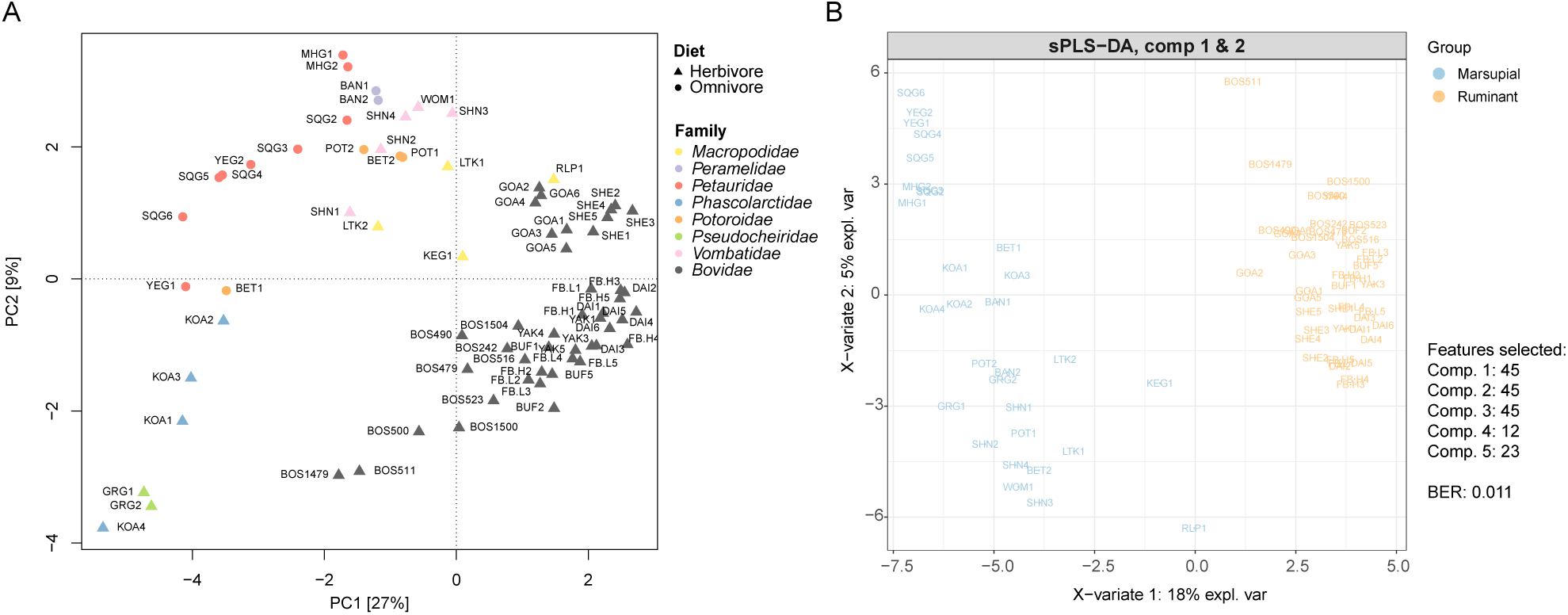
CAZyme-based functional profiles of marsupial and ruminant faecal/rectal samples. (A) Principal component analysis of enzyme abundance profiles. (B) sPLS-DA of enzyme abundance profiles, using marsupial or ruminant as discriminatory variable. Host species are abbreviated as DUN: fat-tailed dunnart, KEG: eastern grey kangaroo, LTK: Lumholtz’s tree kangaroo, RLP: red-legged pademelon, BAN: northern brown bandicoot, MHG: mahogany glider, SQG: squirrel glider, YEG: yellow glider, KOA: koala, BET: rufous bettong, GRG: greater glider, WOM: common wombat, SHN: southern hairy-nosed wombat, BOS: Bos indicus, FB: Bos taurus (Angus), DAI: dairy cattle, BUF: water buffalo, YAK: yak, SHE: sheep, GOA: goat.

**Fig. S8.**
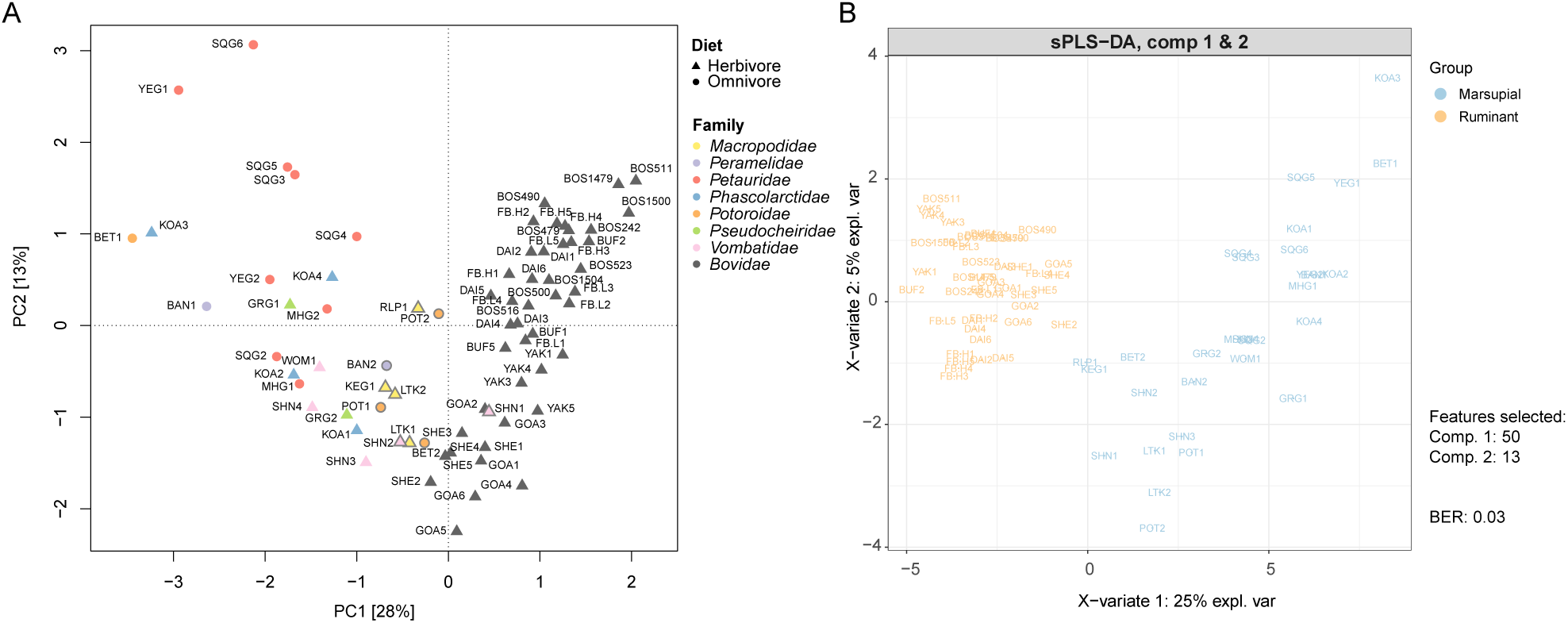
Comparison of hydrogen metabolism functional profiles of marsupial and ruminant faecal/rectal samples. (A) Principal component analysis of protein abundance profiles. (B) sPLS-DA of protein abundance profiles, using marsupial or ruminant as discriminatory variable. Marsupial samples with grey borders are those samples clustered with ruminants in Fig. 3. Host species are abbreviated as DUN: fat-tailed dunnart, KEG: eastern grey kangaroo, LTK: Lumholtz’s tree kangaroo, RLP: red-legged pademelon, BAN: northern brown bandicoot, MHG: mahogany glider, SQG: squirrel glider, YEG: yellow glider, KOA: koala, BET: rufous bettong, GRG: greater glider, WOM: common wombat, SHN: southern hairy-nosed wombat, BOS: Bos indicus, FB: Bos taurus (Angus), DAI: dairy cattle, BUF: water buffalo, YAK: yak, SHE: sheep, GOA: goat.

**Fig. S9.**
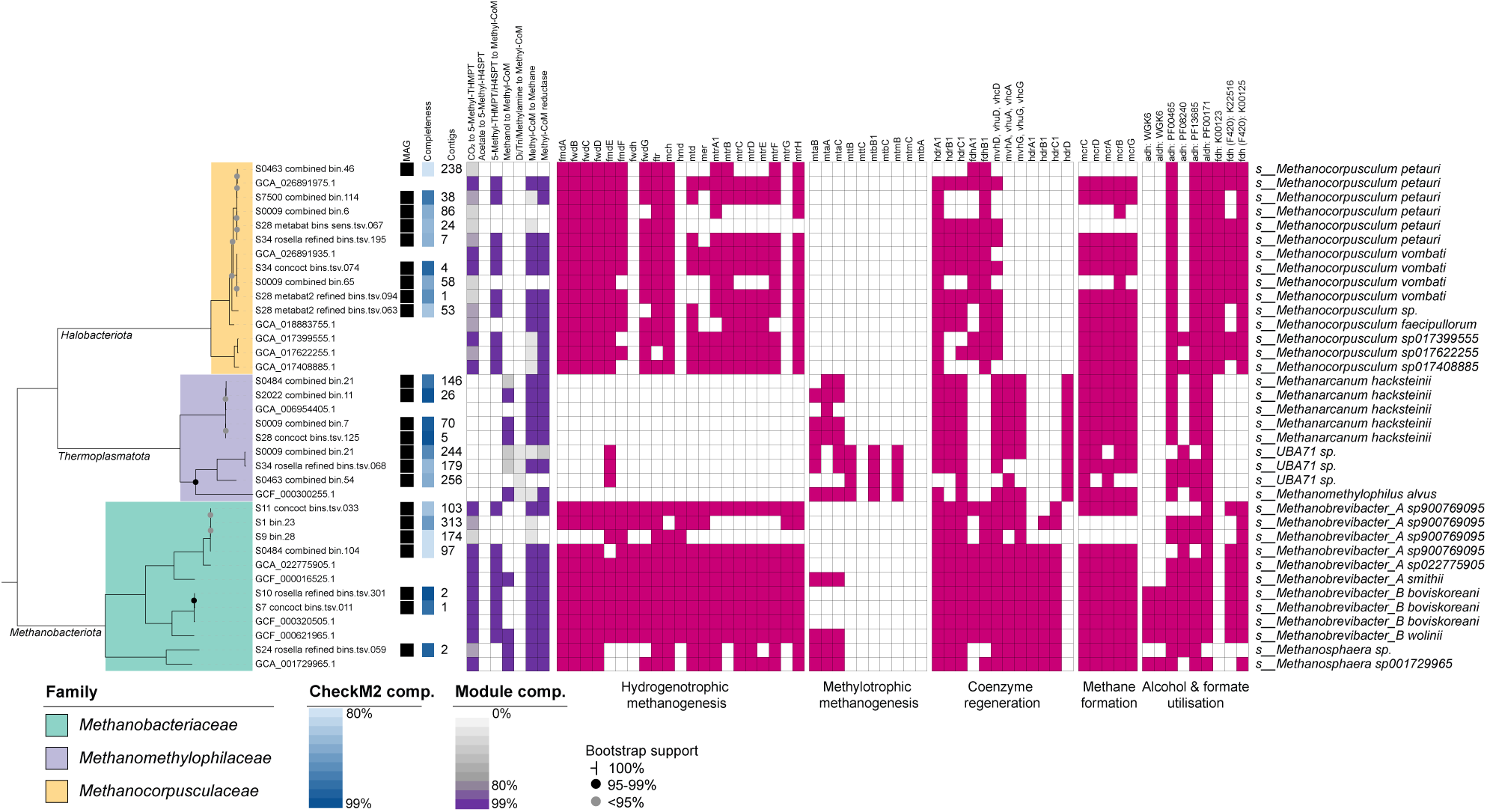
Methanogenesis gene presence across archaeal genomes. Maximum likelihood phylogenetic tree inferred from alignment of 53 GTDB single-copy marker genes. Includes genomes assembled from the dataset exceeding 80% completeness, dereplicated at 80% identity. Supplemented with public genomes representing species present in the dataset or derived from other marsupial studies [22, 23]. Methanogenesis pathway module completeness and individual gene presence indicated for each genome based on KEGG database annotation. Alcohol and formate dehydrogenase presence derived from BLAST-determined homology to WGK6 proteins [23], Pfam or KEGG annotations as indicated.

